# Frontoamygdala hyperconnectivity predicts autonomic dysregulation and persisting symptoms in sports-related concussion

**DOI:** 10.64898/2025.12.18.695278

**Authors:** Kevin C Bickart, Alexander W Kashou, Christopher Sheridan, Allison Lin, Emily L Dennis, Anne Brown, Robert Hamilton, Christopher C Giza, Meeryo Choe

## Abstract

**OBJECTIVE:** We investigated longitudinal trajectories of resting-state fMRI (rsfMRI), autonomic function, and symptoms after sport-related concussion (SRC).

**BACKGROUND:** Limbic circuitry may be particularly vulnerable to traumatic brain injury (TBI), which could explain the affective and autonomic dysfunction some patients develop. Relatively few studies have performed longitudinal rsfMRI analyses in concussion and fewer have combined imaging with autonomic and symptom data. We leveraged published limbic rsfMRI networks centered on the amygdala that include central autonomic structures in frontal and temporal lobes and their visceromotor targets. We hypothesized that frontoamygdala connectivity would differentiate athletes with SRC from matched, in-sport controls (ISC), predict autonomic function, and predict symptom recovery.

**DESIGN/METHODS:** Using independent-samples *t*-tests, we compared rsfMRI connectivity strength in amygdala networks in college athletes with SRC (SRC: n=31, female=14) at three time points after concussion (T1≤4 days, T2=10-14 days, T3=60-90 days) and healthy, matched controls without a concussion in the same sport (ISC: n=36, female=17).

**RESULTS:** SRC athletes showed significantly greater frontoamygdala connectivity compared to ISCs at the acute and subacute post-injury time points (T1 *p*=0.003, T2 *p*=0.014) that normalized to control-level connectivity by the chronic time point (T3 *p*=0.182). When testing whether autonomic function interacts with network connectivity trajectory, we found that opposing trajectories of frontoamygdala connectivity between SRC athletes with higher versus lower acute heart rate variability (HRV), as measured by pNN50 (percentage of intervals between successive normal sinus beats greater than 50ms). SRC athletes with higher HRV acutely post-injury had significantly greater acute frontoamygdala connectivity compared to ISCs at T1; connectivity in these high-HRV SRC athletes normalized to control level over time (T1 *p*=0.001, T2 *p*=0.055, T3 *p*=0.576). SRC athletes with lower HRV acutely post-injury had control-level frontoamygdala connectivity at T1; connectivity in these low-HRV SRC athletes significantly exceeded control-level connectivity at T3 (T1 *p*=0.429, T2 *p*=0.050, T3 *p*=0.002). Furthermore, those SRC athletes with the greatest frontoamygdala connectivity at T3, had the most persistent symptoms on the graded symptom checklist at T3 (r=0.635, *p*=0.001). Differences in diffusion-tensor-imaging-based measures of structural connectivity in the uncinate fasciculus, the fiber bundle most critical to our frontoamygdala network, could not account for these relationships.

**CONCLUSIONS:** These results suggest that increased connectivity in amygdala circuitry acutely after a concussion and its normalization over time may be protective or compensatory; acute measures of amygdala network connectivity and measures of HRV may be valuable biomarkers for predicting symptom persistence.

## Introduction

Traumatic brain injury (TBI), including concussion, leads to an estimated 2.8 million emergency department visits and over 1 million outpatient visits per year in the United States alone ^1,2^. Mild TBI (mTBI) accounts for approximately 80% of these cases, with the highest incidence among teenagers and young adults ^3^. These figures likely underestimate the incidence of mTBI in particular, given evidence that as many as 25% of milder injuries go unreported ^4,5^.

While many patients recover from mTBI within days to weeks, a majority of patients recover from mTBI within 3 months. A substantial minority, however, continue reporting symptoms beyond 3 months, so-called persistent post-concussion symptoms (PPCS), and consume a disproportionate amount of mTBI-targeted care resources seeking effective treatment ^6^. Symptoms typically include heterogeneous constellations of somatic (e.g., headache, neck pain, dizziness, sensory sensitivities), cognitive (e.g., memory and attention deficits), and emotional (e.g., anxiety, post-traumatic stress disorder, depression) problems. PPCS is a significant public health concern; reported rates of PPCS have ranged between 5% ^7^ and 78% ^8^. Consensus among large studies estimates that 40-55% of mTBI patients still report at least one symptom at 3 months post injury ^9,10^. While the field has made progress in understanding the relevant pathobiology and developing validated, context-specific treatments for PPCS, the heterogeneity, subjectivity, and nonspecificity of acute and persistent mTBI symptoms have frustrated significant improvements in patient care ^11–13^.

Maladaptive coping responses, particularly those involving implicit affective regulation, like fear avoidance, play a central role in PPCS development. Affective regulation relies on dynamic interaction between the ventromedial prefrontal cortex (vmPFC), amygdala, and downstream autonomic pathways ^14–20^. These brain regions maintain reciprocal, functionally organized anatomical connections ^21–23^. Some of these functional circuits initiate and reinforce goal-directed behaviors; others extinguish maladaptive habits through repeated exposure ^16^.

Previous research on brain circuit disruption underlying PPCS in humans has focused on major white matter tracts or the largely cortical networks well-characterized in functional imaging literature (i.e., the default-mode network, central executive network, sensorimotor networks, attention networks, etc.), as opposed to limbic networks or, more generally, connectivity related to a specific pathobiological hypothesis ^24^. In contrast, animal models of mTBI have specifically highlighted the vulnerability of frontoamygdala circuitry in dysregulated affective responses. Mild TBI disrupts limbic circuitry in rodents, both increasing excitatory and reducing inhibitory inputs from the vmPFC to the amygdala ^25,26^. These functional changes have led to amygdala hyperconnectivity ^27^ and further transcriptomic changes corresponding to impaired fear extinction ^26^, heightened fear learning ^25,27^, fear overgeneralization ^25^, sensory sensitization ^27^, and anxiety-like behaviors ^28^.

Beyond frontoamygdala circuitry, other areas implicated in concussion symptoms and highly connected to the amygdala are vulnerable to concussive forces, including limbic cortex, hypothalamus, and midbrain nuclei ^29,30^. Most of these regions are critical components of the central autonomic network ^31^. Evidence increasingly shows many people with concussions have abnormalities in heart rate variability (HRV) across a range of injury mechanisms, time since injury, age, and testing conditions. The most consistent finding in concussed patients compared to controls is reduced parasympathetic activity at rest ^32–42^. Investigating dysfunctional central autonomic circuitry, especially in conjunction with HRV or other peripheral biomarkers of autonomic regulation, may be critical to a more actionable understanding of PPCS.

In the present study, we aimed to investigate recovery trajectories of central and peripheral autonomic physiology in people after sports-related concussions (SRC) compared healthy controls matched on age, sex, and sport. We leveraged previously published resting-state fMRI (rsfMRI) parcellations of frontoamygdala circuitry (Figure 1) capturing three anatomically distinct networks supporting different aspects of affective behavior ^43–46^. Two of these networks include regions within the central autonomic network and share connectivity with the vmPFC (red and yellow in Figure 1). Specifically, the “medial” network supports goal-directed behavior through connectivity with visceromotor and reward-related regions (red in Figure 1); the “ventrolateral” network facilitates perception of salient stimuli through connectivity with sensory association regions (yellow in Figure 1). We hypothesized that frontoamygdala connectivity would distinguish athletes with SRC from matched controls, would predict autonomic function post injury, and would predict symptom recovery in SRC subjects. Based on our review of cross-sectional rsfMRI findings in TBI ^24^ and a few prior longitudinal studies ^47–49^, we hypothesized that higher rsfMRI connectivity in the chronic phase would predict greater symptom burden.

**Figure 1.**
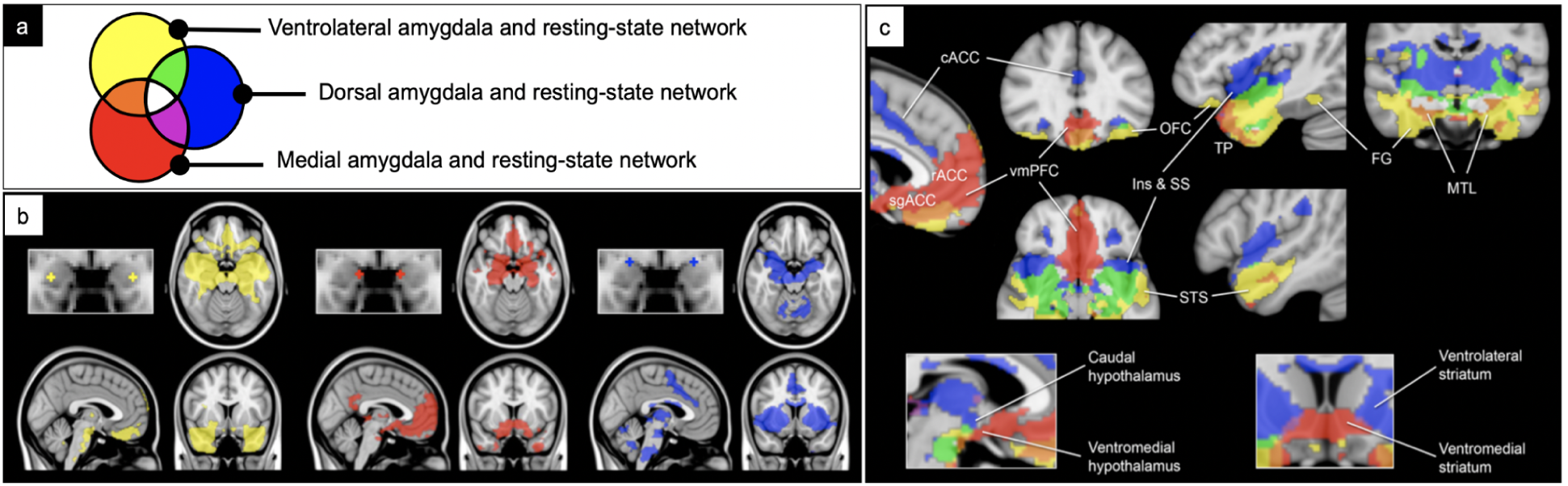
Resting-state networks of the amygdala. (a) Color key. (b-c) Group-level significance maps for amygdala seeds and networks (n=89), binarized at p<10^-5^ and overlaid on a T1 MNI152 0.5mm template (adapted from Bickart et al. 2012 ^45^).

## Methods

### Participants

Club and varsity collegiate athletes aged 14-40 who had recently suffered a diagnosed concussion were recruited and enrolled within 72 hours of reported time of injury (Table 1). Concussions were diagnosed by certified athletic trainers, institutional sports medicine staff, or a healthcare provider at the UCLA Steve Tisch BrainSPORT Clinic. Study staff screened all participants prior to enrollment and ensured all concussions conformed to the study definition of concussion ^50^. Two control groups without a diagnosed or suspected concussion in the 12 months prior to enrollment were recruited. In-sport controls (ISC) were recruited from the same athletic team in the same season as the concussed subject. All controls were matched to concussed athletes on the basis of age, sex, and education. Exclusion criteria were as follows: age <14 or >40 years, membership to an IRB-specified protected class, open head injury, skull fracture, soft tissue trauma to the temporal region, overnight hospitalization for current injury, Glasgow Coma Scale (GCS) ≤12 for current injury, history of seizure disorder, history of a previous moderate or severe TBI, or any contraindication or injury that would prevent participating in the study protocol. All adult participants provided written informed consent. Minor participants provided written assent; their parents provided written informed consent. All study consent procedures and activities were approved by the UCLA institutional review board.

**Table 1.** Demographic variables of mild TBI patients (mTBI) and in-sport controls (ISC).

|  | mTBI (n=31) | ISC (n=36) |
| --- | --- | --- |
| Age: (M, SD) | 20.33 ± 1.36 | 20.50 ± 1.85 |
| Sex (% Female) | 45% (n=14 female) | 47% (n=17 female) |
| Sport (%) | Rugby: 55% (n=17)<br>Soccer: 6% (n=2)<br>Ice Hockey: 6% (n=2)<br>Water Polo: 6% (n=2)<br>Cheer: 6% (n=2)<br>Other: 19% (n=6) | Rugby: 44% (n=16)<br>Soccer: 11% (n=4)<br>Ice Hockey: 6% (n=2)<br>Water Polo: 6% (n=2)<br>Cheer: 6% (n=2)<br>Other: 28% (n=10) |
| # of Prior Diagnosed Concussions (M, SD) | 0.92 ± 1.06 | 0.28 ± 0.61 |
| Days Post-Injury for Scan #1 (M, SD) | 4.16 ± 1.55 | - |
| Days Post-Injury for Scan #2 (M, SD) | 13.23 ± 2.90 | - |
| Days Post-Injury for Scan #3 (M, SD) | 75.88 ± 19.24 | - |

For concussion subjects, the full battery of study assessments was obtained at visits occurring 24-72 hours after injury (T1), 7-10 days after injury (T2), and 60-90 days after injury (T3). For control subjects, the full study battery was obtained at a single time point. Special effort was made to enroll in-sport control subjects within 7 days of their injured counterpart.

### Image Acquisition

All imaging data were obtained on the same 3T Siemens MAGNETOM Prisma Fit whole-body scanner using a dedicated 32-channel head coil.

### Structural Imaging Acquisition

High-resolution T1-weighted structural images were acquired using a magnetization prepared rapid gradient echo (MPRAGE) sequence with the following parameters: FOV=256mm, acquisition matrix=256x256, slice thickness=1.00mm, 176 slices, TR=2300 ms, TE=2.98ms, TI=900ms, flip angle=9, GRAPPA acceleration factor=2.

### BOLD rsfMRI Acquisition

The study participants were scanned for approximately 9 minutes using a gradient-echo, echo-planar image (EPI) sequence with the following parameters: FOV = 245mm, acquisition matrix=70x70, slice thickness=3.5mm, 45 slices, TR=2300ms, TE=27ms, flip angle=90, GRAPPA acceleration factor=2. Participants were instructed to keep their eyes open and to think of nothing in particular during the resting-state session. Prior to each EPI acquisition, a field map was acquired with the following parameters: FOV=245mm, acquisition matrix=70x70, slice thickness=3.5mm, 45 slices, TR=491ms, TE1=4.97ms, TE2=7.43ms, flip angle=60.

### Preprocessing and Analysis of BOLD rsfMRI

Image preprocessing was performed via the default SPM12 pipeline as implemented in the CONN toolbox (version 19). All MRI data were processed using SPM12 (Wellcome Department of Cognitive Neurology, London, UK) implemented in MATLAB 2019b (Mathworks Inc, Natick, MA). Before processing, images were converted from DICOM to 4D NifTi format. Structural, T1-weighted images were segmented into their component parts (gray matter, white matter, CSF, skull, and background), bias corrected, and normalized simultaneously (SPM: segment). Gray matter, white matter, and CSF tissue segmentations were then applied as a combined mask to remove the skull from the anatomical image (SPM: imcalc). Field map-derived voxel displacement maps were generated and co-registered to the functional images (SPM: VDM). Functional images were then realigned with rigid-body 6 motion parameter realignment to the first image in the series using a least-squares approach. Previously calculated voxel displacement maps were used to unwarp the functional images, reducing image distortion and preserving anatomical fidelity (SPM: Realign and Unwarp). The functional images were then co-registered, through the mean T2* image, to the skull-stripped anatomical (SPM: Coregister: Estimate). The co-registered images were then normalized to standard MNI-152 space using the nonlinear deformation maps obtained during segmentation. Lastly, functional images were skull-stripped and smoothed with an 8mm full-width at half-maximum (FWHM) kernel to enhance the signal-to-noise ratio and reduce inter-subject variability.

To calculate resting-state functional connectivity (rsFC) between the medial and ventrolateral amygdala networks, we computed ROI-to-ROI correlations of averaged BOLD signals across previously published masks for each network (Figure 1, red and yellow). The resulting Pearson correlation values were converted to Fisher z values for further analysis.

### HRV Acquisition and Analysis

Blood volume pulse waveform data were acquired from participants during the resting-state BOLD sequence via a BIOPAC TSD200-MRI photoplethysmograph (PPG) transducer affixed to the left index finger of the subject. Signal was sampled at a frequency of 1000Hz and transmitted the data to a BIOPAC MP150 system via a BIOPAC PPG100C-MRI amplifier unit. PPG traces were time-locked to MRI sequences via a transistor-transistor logic (TTL) trigger.

Preprocessing, quality control, and analysis of PPG data were performed using BIOPAC’s AcqKnowledge software (v5.0.5). Data were bandpass filtered with a Finite Impulse Response (FIR) filter between 0.3 Hz and 35 Hz. A peak detection algorithm labeled and extracted successive R-waves. Subsequently, all traces were visually inspected to identify artifacts and determine accurate R-wave detection. Incorrectly labeled peaks were removed, and R-waves missed by the algorithm were labeled manually. If any artifacts or missing data were present, those data were excluded from further analyses. Traces with fewer than two continuous minutes of acceptable quality data were excluded from analysis.

### Symptom Assessment

Subjects provided demographic and health history information and completed a battery of questionnaires and neuropsychological tests. We used the Graded Symptom Checklist (GSC) from the Sport Concussion Assessment Tool - 3rd Edition (SCAT3) to assess symptom persistence. The GSC captures self-reported symptom severity on a 22-item inventory with each symptom ranked on a Likert scale from 0 (none) to 6 (severe). Symptom total score (number of symptoms endorsed at above 1 in severity) ranges from 0 to 22, and symptom severity score ranges from 0 to 132 ^51^. In addition to repetition of the clinical assessment battery, information about concussion management and symptom recovery, including number of days symptomatic and RTP decision, was collected at follow-up visits.

### Statistical Analysis

To ensure comparable groups, we examined sample characteristics with two-tailed t-tests. Resting-state functional connectivity (rsFC) between the medial and ventrolateral amygdala networks was then compared across SRC and ISC groups using an ANOVA to test for group-by-time interactions as well as the main effects of time and group. To explore interactions between HRV and rsFC, we correlated HRV from the initial time point (T1) with rsFC between the medial and ventrolateral amygdala networks at T1, T2, and T3 within the concussed group. Based on these results, we split the SRC group into high and low HRV subgroups and conducted an ANOVA to test for HRV-by-time interactions, followed by main effect tests of HRV and time. Correlations between rsFC in the medial-ventral amygdala networks and symptoms were analyzed using Pearson’s R. Next, voxelwise analyses identified specific regions within the medial and ventrolateral amygdala networks contributing to symptoms and HRV. The Benjamini-Hochberg (false discovery rate [FDR]) procedure was used to account for multiple comparisons. Finally, for functional specificity, we examined whether fractional anisotropy (FA) averaged across the left and right uncinate fasciculi differed over time or correlated with symptoms. For all statistical tests, an alpha level of 0.05 was set to determine significance.

## Results

### Sample Characteristics

Groups were well-matched for age, sex, and sport (Table 1). We investigated longitudinal differences in resting-state connectivity at three time points after a sports-related concussion (T1≤4 days, T2=10-14 days, T3=60-90 days) in male and female collegiate athletes (SRC, n=31, female=14) and group differences between these concussed athletes and healthy age-and sex-matched controls in the same sport (in-sport control [ISC], n=36, female=17). Days post-injury for MRI scanning in the SRC group corresponded to acute (T1=4.2d), subacute (T2=13.2d) and chronic (T3=75.9d) (Table 1).We specifically investigated connectivity differences between the medial and ventrolateral frontoamygdala networks.

### Trajectory of Amygdala Network Connectivity

Compared to controls, SRC athletes showed increased connectivity between the medial and ventrolateral amygdala networks at T1 (*p*=0.003) and T2 (*p*=0.014) that normalized to control-level by T3 (*p*=0.182). Specifically, there was a parametric decrease in connectivity between networks over time, starting significantly higher than controls post injury and falling to comparable levels by the T3 (Figure 2).

**Figure 2.**
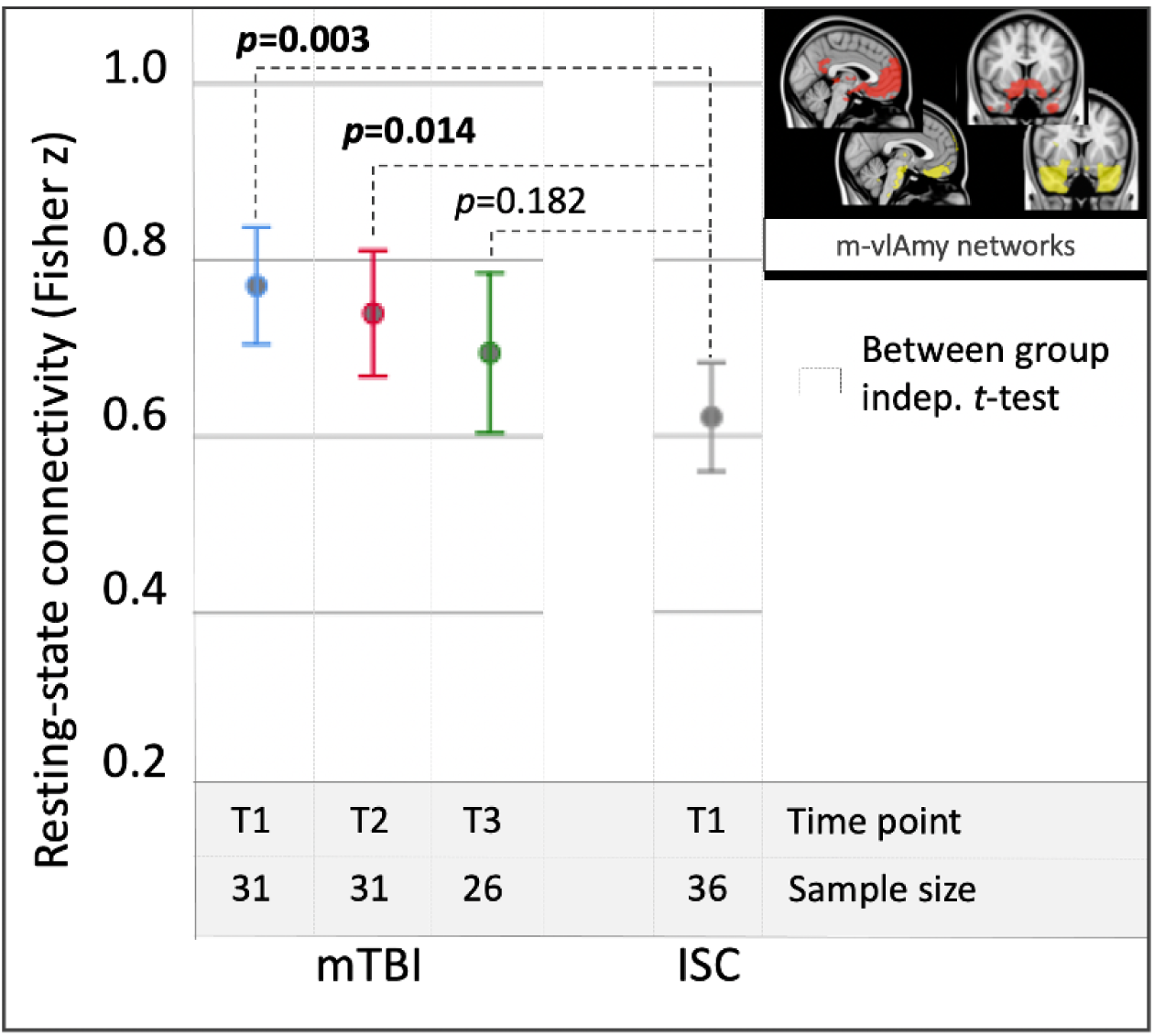
Longitudinal amygdala network connectivity in concussed versus in-sport control (ISC) athletes.

### HRV Interaction with Amygdala Network Connectivity

Given the autonomic and visceromotor role of regions within these networks (e.g., amygdala, vmPFC, nucleus accumbens, hypothalamus, and ventral anterior insula), we investigated whether a metric of autonomic function might mediate the above finding. HRV (pNN50) at T1 significantly predicts higher between-network connectivity at T1 (Fisher’s *r^2^*=0.32, *p*=0.005), shows no correlation at T2 (Fisher’s *r^2^*=0, *p*=0.9), and predicts lower between-network connectivity at T3 (Fisher’s *r^2^*=0.29, *p*=0.016; Figure 3). That is, SRC athletes with higher HRV at T1 tended to have higher connectivity at T1 and lower connectivity at T3 relative to SRC athletes with lower HRV.

**Figure 3.**
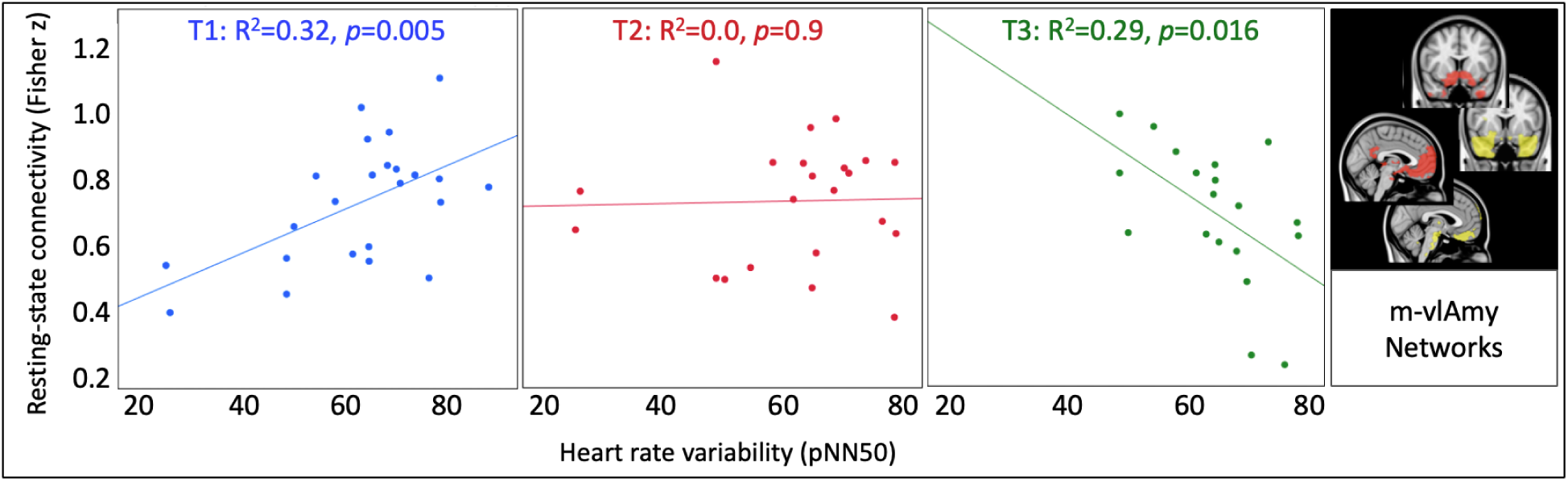
HRV interacts with amygdala network connectivity trajectory after concussion.

We subsequently stratified the SRC group by HRV (pNN50) at the 50th percentile (pNN50=65%) and re-analyzed between-network connectivity. This stratified analysis confirmed a significant interaction between HRV group and time (F=13.85, *p*=0.002). SRC athletes with higher HRV (HHRV) and SRC athletes with lower HRV (LHRV) at T1 exhibited opposing between-network connectivity trajectories (Figure 4). Compared to ISC, HHRV athletes had significantly higher than normal connectivity at T1 and T2 that normalizes by T3 whereas LHRV athletes have normal connectivity at T1 that parametrically increases to significantly higher than normal connectivity by T2 and T3.

**Figure 4.**
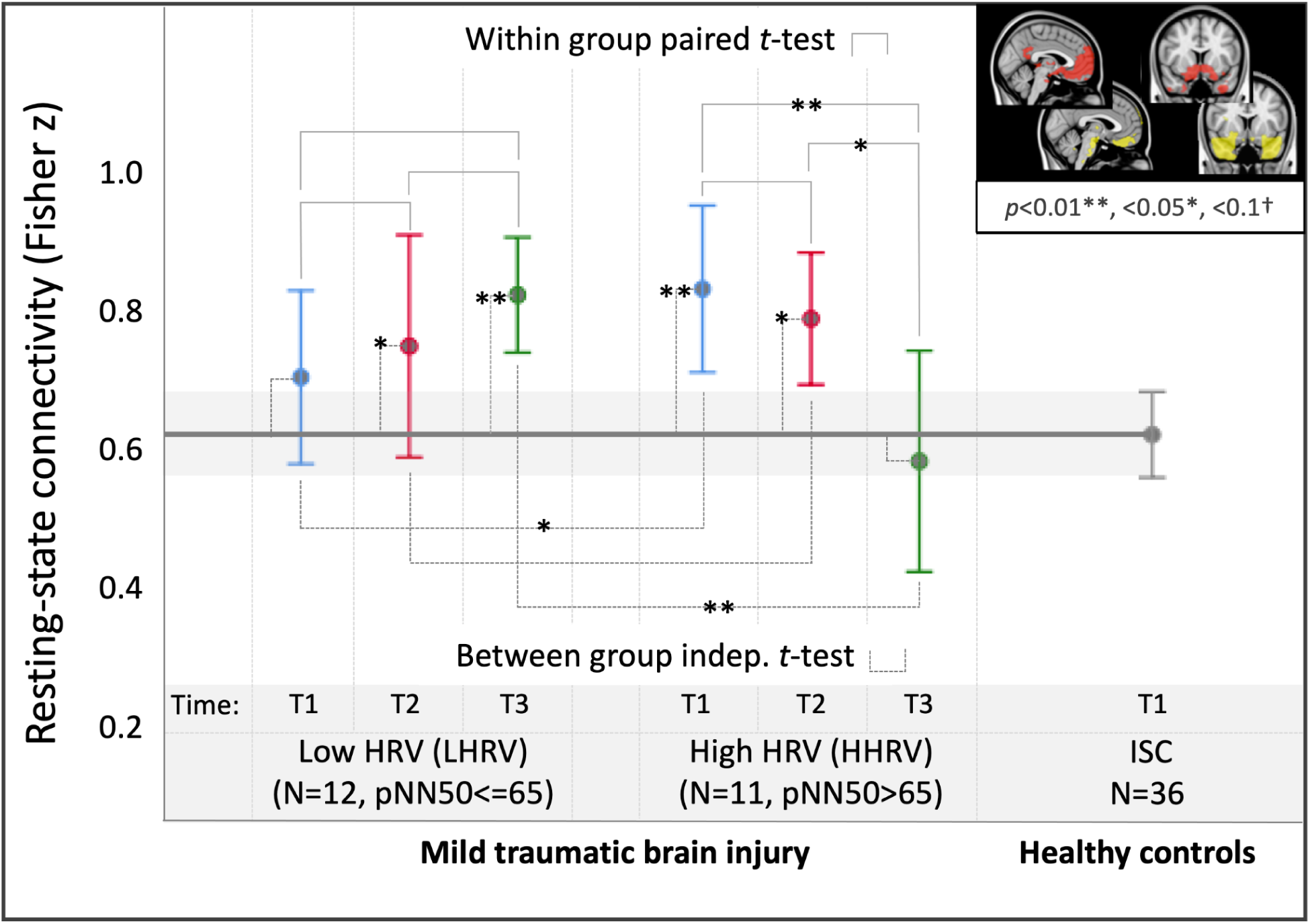
Concussed athletes with high (HHRV) versus low heart rate variability (LHRV) acutely after concussion show opposite trajectories of amygdala network connectivity.

### Amygdala Network Connectivity Predicts Persistent Symptoms

Furthermore, for the concussion group as a whole, those with the greatest amygdala network connectivity at the chronic time point (T3) had the greatest persisting concussive symptoms on the GSC, no matter their HRV status (Figure 5).

**Figure 5.**
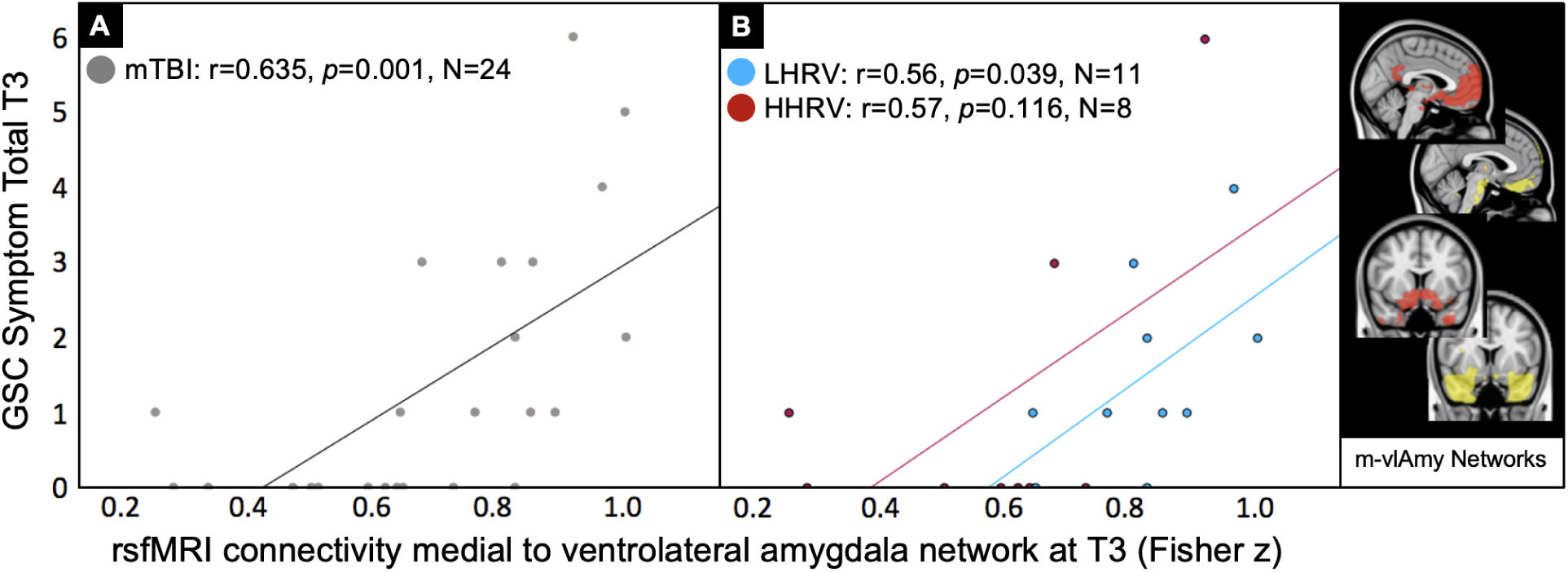
Higher chronic amygdala network connectivity predicts worse persistent symptoms independent of HRV. (A) Mild traumatic brain injury (mTBI) amygdala connectivity correlated with symptom severity at T3. (B) Low (LHRV) and high heart rate variability (HHRV) subgroup amygdala connectivity correlated with symptom severity at T3.

### Higher Amygdala-vmPFC Connectivity Predicts Persistent Symptoms and Lower HRV

Within the medial and ventrolateral amygdala networks, it was higher amygdala connectivity to voxels in the vmPFC that best predicted persistent symptoms at the chronic timepoint T3 (Figure 6A) and higher initial HRV (Figure 6B).

**Figure 6.**
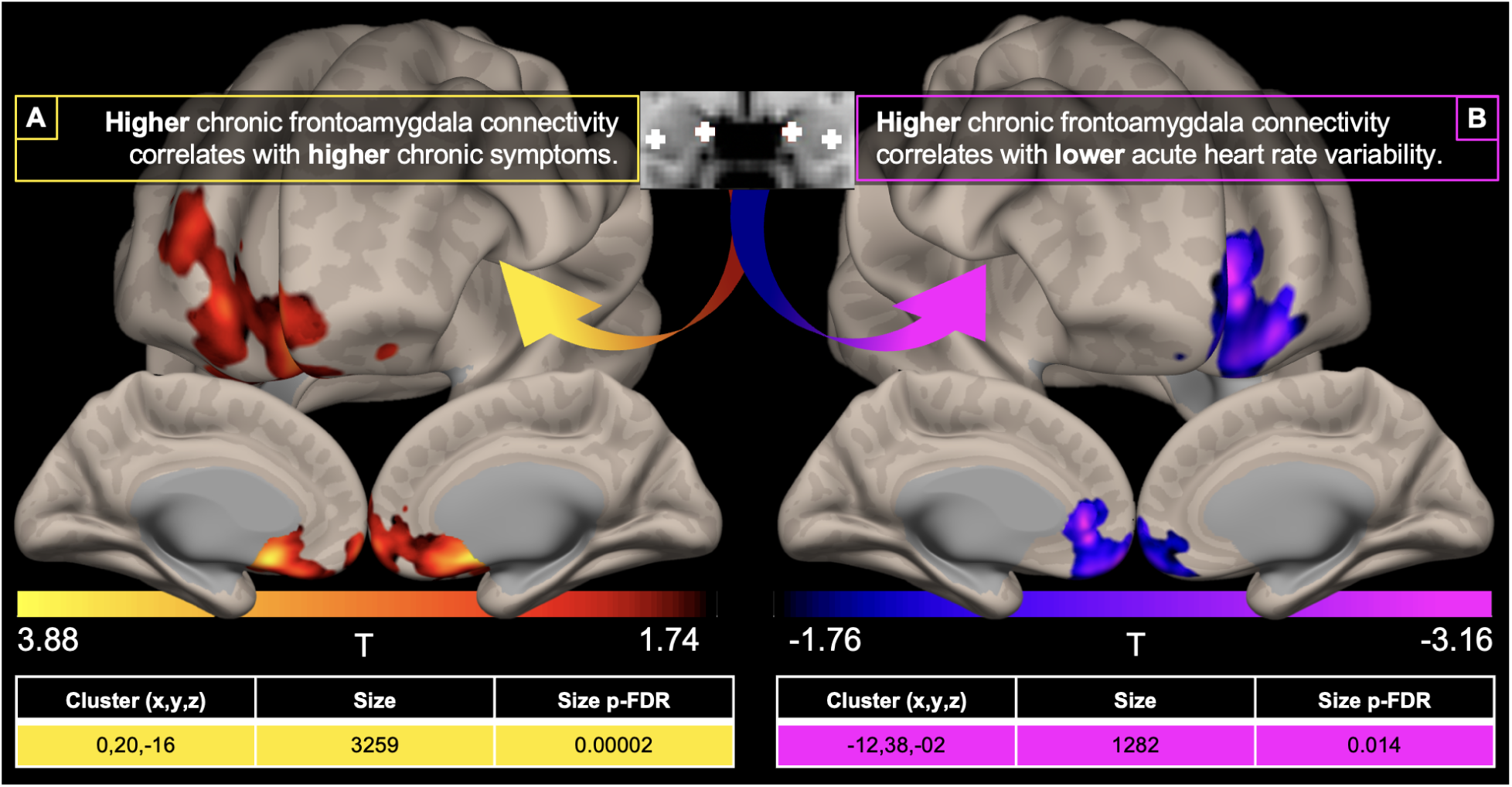
Higher chronic frontoamygdala connectivity correlates with worse chronic symptoms and lower acute HRV. (A) Voxelwise regression of combined medial and ventrolateral amygdala connectivity at T3 by GSC symptom total from T3. (B) Voxelwise regression of combined medial and ventrolateral amygdala connectivity at T3 by HRV (pNN50) at T1. Both analyses used GLM performed in the CONN Toolbox with voxel threshold of p-uncorrected <0.05 and cluster threshold p-FDR corrected <0.05, restricted to positive voxels (A) or negative voxels (B).

### Frontolimbic Structural Integrity Does not Account for Functional Connectivity Results

We also investigated the integrity of the main structural pathway connecting the amygdala to the vmPFC, the uncinate fasciculus. Similar to the rsfMRI trajectory, bilateral uncinate fractional anisotropy (FA) parametrically decreased from the acute (T1) to the chronic (T3) time point after concussion as compared to ISCs (Figure 7A). This trajectory did not account for variation in rsfMRI findings and did not predict symptoms at the chronic time point (Figure 7B).

**Figure 7.**
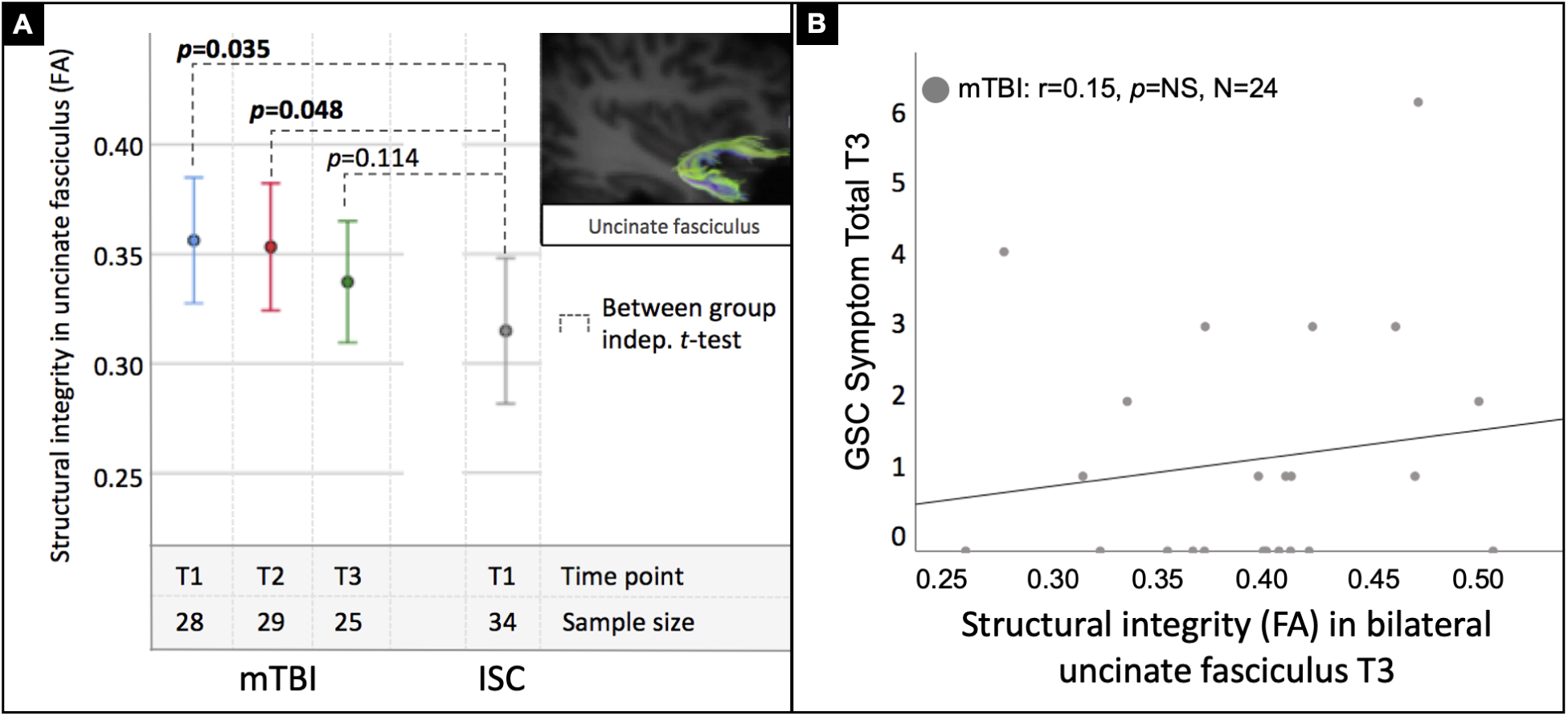
Uncinate fractional anisotropy (FA) decreases over time but does not predict chronic concussion symptoms. (A) FA in bilateral uncinate fasciculus parametrically decreases over time for the concussed group becoming indistinguishable from controls by the chronic time point. (B) Chronic FA however, did not predict chronic symptom burden for the concussed group.

## Discussion

The present study investigated recovery trajectories of central and peripheral autonomic physiology with rsfMRI and HRV in sport-related concussion as compared to age, sex, and sport-matched healthy controls. We leveraged previously published rsfMRI parcellations of frontoamygdala circuitry containing regions within the central autonomic network with overlapping connectivity in the vmPFC to evaluate longitudinal changes that may predict recovery of symptoms and return to play.

We found that concussed athletes with higher versus lower HRV acutely show opposing trajectories of frontoamygdala connectivity over the first 90 days post-injury. Specifically, athletes with higher HRV showed significantly increased connectivity acutely that normalized parametrically over time whereas athletes with lower HRV showed connectivity in the normal range acutely that increased parametrically over time to be significantly elevated over controls by the chronic time point. This suggests that higher HRV and higher frontoamygdala connectivity in the acute phase after injury may be protective and predictive of an adaptive and more rapidly normalizing neurophysiological response, either marking individuals who were more fit prior to injury or more durable to injury. In contrast, acutely lower HRV and lower frontoamygdala connectivity may be a maladaptive response or suggest preexisting vulnerability that predicts a rise in connectivity over the time window to abnormal levels.

In the context of prior work, acutely elevated frontoamygdala connectivity that normalizes within the first three months post-injury may reflect more effective regulation of the autonomic nervous system. The link between autonomic function and frontoamygdala circuitry is well established ^52–55^. Prior fMRI studies, across both task-based and resting-state conditions as well as in both humans and nonhuman primates, have demonstrated correlations between autonomic measures (e.g., HRV, heart rate, blood pressure) and coactivation or connectivity within frontoamygdala circuits ^21,52,56–60^. In one such study, Wong et al. recorded heart rate (HR) and mean arterial pressure (MAP) while assessing cortical activity via fMRI during a graded handgrip task. They found that HR responses inversely correlated with activity in the vmPFC, a region with dense anatomical connections to subcortical autonomic centers, supporting the hypothesis that the vmPFC, along with its subcortical partners, plays a key role in modulating vagal efferent output ^53^. Other studies have echoed these findings, such as Motzkin et al., who showed that lesions to the vmPFC in humans were associated with increased amygdala activity and corresponding autonomic dysfunction, or Sakaki et al., who demonstrated reciprocal fluctuations in vmPFC and amygdala activity that tracked with HRV ^56,58–60^. Our findings build on this foundation by using longitudinal assessments of both connectivity and autonomic function over the course of recovery.

Heart rate variability (HRV) may serve as a valuable proxy for central autonomic disruption following traumatic brain injury (TBI). Indeed, autonomic dysfunction is a well-documented consequence in moderate to severe cases of TBI ^61–68^. Even among individuals with mild TBI or concussion, HRV can remain altered for months or years post-injury, distinguishing them from healthy controls ^34,40,41,69,70^. Previous work has shown that the vmPFC and amygdala are particularly vulnerable to concussive forces, which can disrupt the top-down modulation of peripheral physiological systems ^71–75^. These disruptions may contribute to altered HRV and broader dysautonomia following injury. However, it is important to note that HRV is inherently nonspecific and can be influenced by a range of other systemic perturbations. Future investigations that more directly delineate the mechanistic link between concussion and HRV, potentially through the use of high-fidelity wearable devices and continuous physiological monitoring, may offer a powerful means of tracking recovery trajectories in concussed individuals over chronic periods.

These insights into the vulnerability of central autonomic structures may open the door to treatment for chronic concussive symptoms. Lower frontoamygdala connectivity and HRV acutely, or elevated connectivity chronically, may represent viable therapeutic targets. Such treatments could involve modalities known to influence this circuitry, including graded exposure therapy (e.g., progressive return to physical activity), neuromodulation, or neurofeedback. Interventions like vagal nerve stimulation (VNS) and heart rate variability biofeedback (HRV-BF) have shown promise in mitigating TBI-related deficits ^76^. Both methods aim to modulate vagal nerve activity, which projects broadly throughout the brain, including forebrain and limbic regions^77^. VNS has demonstrated improvements in cognition and motor function in animal models^76,78,79^, though its application in human mild TBI (mTBI) populations remains unexplored. HRV-BF, a non-invasive technique that uses diaphragmatic breathing to elicit vagal responses ^80–83^, has been associated with enhanced cognitive performance (i.e. attention, working memory, short-term memory, inhibition), improved emotional regulation, and reductions in post-concussive symptoms in both mTBI and more severe TBI cases ^77,84–86^. Our study supports the potential utility of these interventions by identifying altered functional connectivity patterns in the amygdala and its vagal projections, suggesting that rsfMRI, paired with long-term HRV monitoring, may help identify individuals most likely to benefit from targeted neuromodulatory or behavioral therapies.

Interestingly, the progressive decrease in FA within the primary structural pathway connecting the amygdala to the vmPFC from T1 to T3 did not predict symptoms at the chronic timepoint, nor did it significantly correlate with functional connectivity measures. This suggests that the clinical manifestations of mTBI are more driven by functional alterations than by structural differences in this circuitry in particular. Such perturbations are frequently observed in chronic concussion populations, making this finding consistent with prior studies ^87–89^. Nonetheless, the literature on DTI outcomes in mTBI remains heterogeneous, highlighting the need for standardized imaging protocols and demographically uniform patient cohorts to better evaluate the predictive value of DTI in TBI patients without radiographically visible lesions.

## Limitations

This study was relatively limited in sample size for generating reliable brain-behavioral correlations. Our discrete phenotype, sport-related concussion in college sports, rigorous control sample (sport-matched non-concussed athletes), and longitudinal MRI, clinical, and autonomic assessments help to strengthen the findings. Ideally, resting-state protocols would include greater than 12 minutes of data. The protocol here included about 9 minutes of data. It was, however, high quality and is above the average compared to published literature ^24^.

We derived HRV from only 2 minutes of clean data from a baseline period of rest while the participants were undergoing their rsfMRI scan. This has some limitations. First in the short timeframe of acquisition. Two minutes is the minimum needed but 5 minutes or greater is more reliable, especially for the lower frequency domain metrics. In this case, we used a time domain metric (pNN50), which is less sensitive to slower fluctuations in interbeat intervals that drive low frequency domain computations.

Lastly, the control group did not undergo repeat scans, so the natural fluctuations in connectivity cannot be directly assessed as compared to the injured patients over time, but only to the control group’s initial timepoint. Future studies would benefit from longer rsfMRI and HRV acquisitions as well as longitudinal data in both the injured and control populations.

## Conclusion

This study integrates central and peripheral autonomic physiology to demonstrate unique trajectories of recovery after sport-related concussion. Concussions can lead to a temporary disruption in functional connectivity, which in turn contributes to autonomic dysfunction, observable through HRV measures. In this study, integrating rsfMRI functional connectivity with HRV data highlights a promising approach to identifying individuals at greater risk for developing PPCS. Our findings suggest that individuals with high HRV (HHRV) shortly after injury display increased rsFC, which normalizes over time relative to healthy controls, correlating with a lower likelihood of PPCS. Conversely, those with low HRV (LHRV) in the acute phase exhibit reduced rsFC that increases over time, aligning with a higher risk of PPCS. These results represent a potential future state of personalized concussion management, where HRV and rsfMRI data may forecast recovery trajectories and pinpoint candidates for therapeutic intervention, ultimately enhancing athlete safety and reducing long-term consequences of concussions.

## Acknowledgements

The authors have no acknowledgements at this time.

## Contributions

**K.C. Bickart:** Conceptualization, Data Curation, Formal Analysis, Investigation, Methodology, Project Administration, Resources, Software, Supervision, Validation, Visualization, Writing - Original Draft, Writing - Review and Editing. **A. Kashou:** Investigation, Resources, Validation, Visualization, Writing - Original Draft, Writing - Review and Editing. **C. Sheridan:** Conceptualization, Data Curation, Investigation, Methodology, Resources, Validation, Writing - Original Draft, Writing - Review and Editing. **A. Lin:** Data Curation, Formal Analysis, Investigation, Validation, Writing - Review and Editing **E.L. Dennis:** Formal Analysis, Writing - Review and Editing. **A. Brown:** Data Curation, Formal Analysis, Investigation, Writing - Review and Editing. **C.C. Giza:** Investigation, Validation, Writing - Review and Editing. **C. Thibeault:** Investigation, Validation, Writing - Review and Editing. **R. Hamilton:** Investigation, Validation, Writing - Review and Editing. **M. Choe:** Conceptualization, Data Curation, Investigation, Methodology, Resources, Supervision, Validation, Writing - Original Draft, Writing - Review and Editing.

## Conflicts of Interest

The authors have no competing interests to disclose.

## Funding

This research was supported by the following funding: NIH SBIR R44NS092209 in partnership with NeuraSignal Inc.; the UCLA Steve Tisch BrainSPORT Program; UCLA Easton Clinic for Brain Health; UCLA Brain Injury Research Center; The Harry O. Parker Neuroscience Research Fund; and Stanley and Patti Silver.

## Notes

### Competing Interest Statement

The authors have declared no competing interest.

